# Precision identification and prediction of high mortality phenotypes and disease progression pathways in severe malaria without requiring longitudinal data

**DOI:** 10.1101/425132

**Authors:** Till Hoffmann, Iain Johnston, Sam Greenbury, Ornella Cominetti, Muminatou Jallow, Dominic Kwiatkowski, Mauricio Barahona, Nick Jones, Climent Casals-Pascual

## Abstract

The parasite *Plasmodium falciparum* is the main cause of severe malaria (SM). Despite treatment with antimalarial drugs, more than 450,000 SM deaths are reported every year, mainly in African children. The diversity of clinical presentations associated with SM indicate important differences in disease pathogenesis that often require specific treatment, and this clinical heterogeneity of SM is largely unresolved. In this study, we apply new machine learning and inference tools for large-scale data analysis to dissect the heterogeneity in patterns of clinical features associated with SM in 2,695 Gambian children admitted to hospital with *Plasmodium falciparum* malaria. This quantitative analysis, including the powerful HyperTraPS algorithm for inference of progressive processes, reveals pathways of SM symptom progression and features predicting the severity of individual patient outcomes. Notably, our approach allows the identification and dissection of disease progression pathways without the need for longitudinal observations. Learning these pathways and features from this rich dataset allows us to construct several quantitative measures of the mortality risk associated with a patient presenting with a given set of symptoms. By independently surveying expert practitioners, we show that this data-driven approach agrees with and expands the current state of knowledge on malaria progression, while simultaneously providing a data-supported framework for predicting clinical risk.

## Introduction

Severe malaria (SM) is a major public health problem and a complex disease. Worldwide, 3.3 billion people live in areas where malaria is transmitted by infected anopheline mosquitoes. Despite recent improvements in the implementation of effective control measures in some countries, an estimated 216 million clinical malaria cases and 445,000 deaths were reported in 2016, mainly in Sub-Saharan Africa [1].

The definition of severe malaria proposed by the World Health Organization (WHO) was designed to capture the majority of children at risk of dying, and thus it prioritizes sensitivity over specificity. In sub-Saharan Africa, children with coma (cerebral malaria) and/or respiratory distress are at the highest risk of death. Yet these clinical syndromes encapsulate a heterogeneous population, and possibly reflect diverse pathophysiological processes. Critically, the current WHO classification of SM fails to capture this heterogeneity and thus treatment allocation based on this definition may have undesired consequences. Indeed, most adjuvant treatments proposed to date have consistently failed to improve patient outcome, and some of these treatments have been shown to increase mortality in children [2-4].

The sequence of events leading to severe malaria is poorly understood. A major determinant of death is the time elapsed from the initial symptoms to clinical presentation, with most deaths taking place within 24 hours of admission [5-7]. Typically, clinical studies rarely capture the temporal component of the infection; accordingly, the natural history of the disease is inferred from experimental models, even though findings from these models are not always easily translated to human malaria [8].

The explosion of data generation across the biomedical sciences coupled with advances in mathematical and computational tools provide the unprecedented opportunity to learn about these poorly understood dynamics [9]. Quantitative approaches leveraging these large datasets can be used to reveal progression pathways and learn features correlated with patient outcomes. Such approaches are central to the ongoing goal of “precision medicine”, where clinical protocols are optimally tailored to the individual under consideration. However, the heterogeneity and the large scale of biomedical data, including data on malaria progression, poses challenges to quantitative approaches. Typically, the identification of prognostic factors is based on *generalised linear models* that use a set of features as *independent* variables. However, the complexity of these models increases dramatically when interactions between factors are important, and generalised linear models lack the ability to naturally dissect dynamic data.

Here, we applied more powerful, recent tools to analyse the rich data on SM progression to facilitate robust clinical statements about disease progression dynamics and risks associated with individual patients. To this end, we combine mutual information (MI), used to learn clinical factors predictive of patient outcomes, with the recently developed HyperTraPS (hypercubic transition path sampling) algorithm [10], which we use to learn dynamic probabilistic pathways of disease progression. MI approaches are more robust regarding nonlinearities in relationships and do not suffer from the shortcomings of log odds ratios associated with linear regression. HyperTraPS, which was recently applied to elucidate the dynamics of and mechanisms underlying the evolution of mtDNA genome structure [10] and efficient photosynthesis [11], allows the Bayesian inference of dynamic pathways describing the successive acquisition of features or symptoms directly from cross-sectional (or longitudinal) observations. Here we apply MI and HyperTraPS to identify prognostic factors and to infer the sequence of events from cross-sectional data in patients with severe malaria. We underline that the approaches we describe are not specific to severe malaria and have general applicability to the study of disease progression. We compare the results of our data-driven analysis with a survey of expert opinion on malarial progression and demonstrate how these quantitative methods can be used to make predictions in precision medicine approaches for patient stratification.

## Results

### Features informative of death beyond the WHO-classification

The analysis included 2,915 patients with severe *Plasmodium falciparum* malaria. Clinical features of disease severity were used to classify patients into three overlapping clinical categories: 1,166 children had respiratory distress (40.1%); 1,060 (36.5%) had CM; and 659 (22.7%) had severe anaemia. There were 387 deaths among 2,904 children admitted with SM for whom outcome data were available (case-fatality rate: 13.3%).

We used mutual information (see Methods) to seek the features that most strongly correlated with mortality in our datasets. We employed the following iterative procedure: we identified the feature that best predicts mortality for a set of observations and split the dataset according to that feature; we then sought the next feature that best predicts mortality. The algorithm stops when no further statistically robust connections with mortality can be identified.

Using this approach we found that presence of *cerebral malaria* was the strongest initial predictor of mortality (Fig 1). Splitting the cohort into cerebral versus non-cerebral malaria cases, we found that the presence of respiratory distress was the next most predictive feature of mortality in both cases. Further iterations of the MI evaluation revealed that abnormal posturing, absence of transfusion and lack of splenomegaly were robust predictors of mortality. The stratification afforded by this iterative approach can reveal the features most strongly linked to mortality for any given combination of clinical presentations (see Methods).

**Figure 1.**
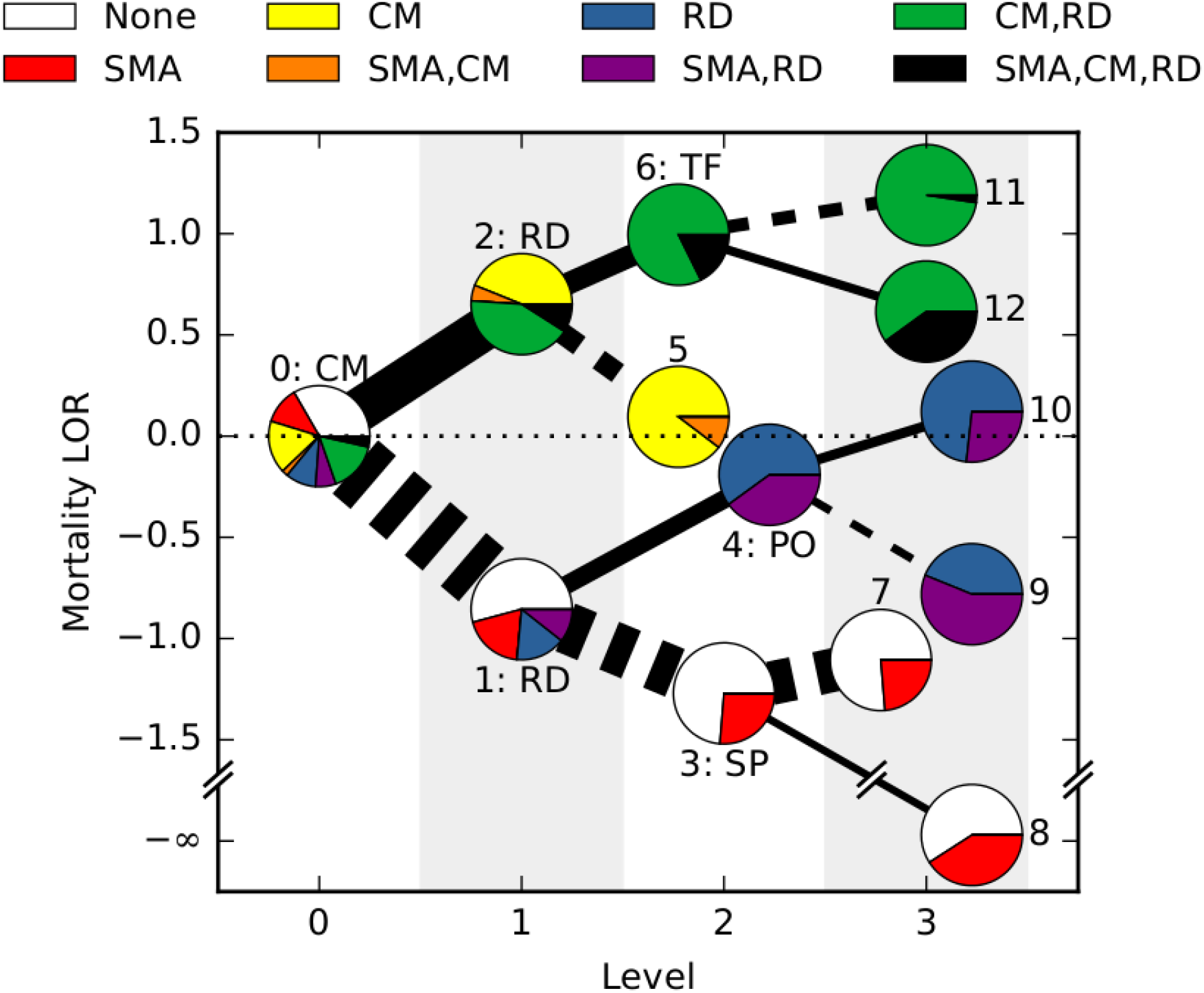
Mutual information approach to identify features predicting mortality. At each level (horizontal axis), patient data are greedily split into two subsets according to the remaining feature that most strongly predicts mortality. The algorithm stops when no feature is statistically significantly associated with death. The figure shows a tree generated by this algorithm: cerebral malaria (CM), respiratory distress (RD), splenomegaly (SP), abnormal posturing (PO), transfusion (TF) are selected as informative features. Nodes are shown as pie charts representing the composition of WHO classifications in each cluster. Solid (dashed) edges indicate that the feature was present (absent) and their width is proportional to the number of patients. The vertical axis corresponds to the mortality log odds ratio compared with the average mortality. Partition 8 has infinite LOR because all patients survive.

### Sequence of events leading to severe malaria

To next elucidate the pathways by which malaria progresses, we used HyperTraPS (hypercubic transition path sampling), an algorithm for inferring the dynamics by which traits are acquired or lost over time [10]. The clinical features in our dataset comprise a collection of true/false and ordinal features. In the case of true/false features, “true” equates feature presence and “false” feature absence (for example, “Is the patient coughing?”). In some case (see Methods) these observations correspond to a quantitative threshold; for example, hyperparasitaemia is defined as over 2.5 × 10^5^ parasites/µL blood). Regarding ordinal features, we consider a set of inequalities reporting the severity of the condition. For dehydration, the score runs from 0 to 3: we assign separate features describing when this score is ≥ 1, ≥ 2, and =3. For consciousness we use the Blantyre coma score, which ranges from 5 (fully conscious) to 0 (not responsive): we assign separate features describing when this score is ≤ 4, ≤ 3, ≤ 2, ≤ 1, and =0. Importantly, we take into account that these inequality features are not independent, i.e., a BCS score ≤ 2 is also ≤ 4. In consequence, we expect to see an ordering in any posteriors on these features.

Two features were treated differently: death and transfusion. We used death to split the dataset into those patients that survived and those that died: these different sets are kept separate in the analyses for differences in the pathways associated with these different outcomes to surface. Since transfusion is an extrinsic intervention rather than an intrinsic feature of disease progression, we removed it from the inference of disease pathways for most of the subsequent analysis. If transfusion is retained in the dataset, it is identified as a significant discriminant between survivor and death pathways, in agreement with the results above (Fig S1). After removing this feature, we computed posterior distributions over disease progression pathways separately for cases where the patient survived and cases in which the patient died. These posteriors are summarised in Fig 2.

**Figure 2.**
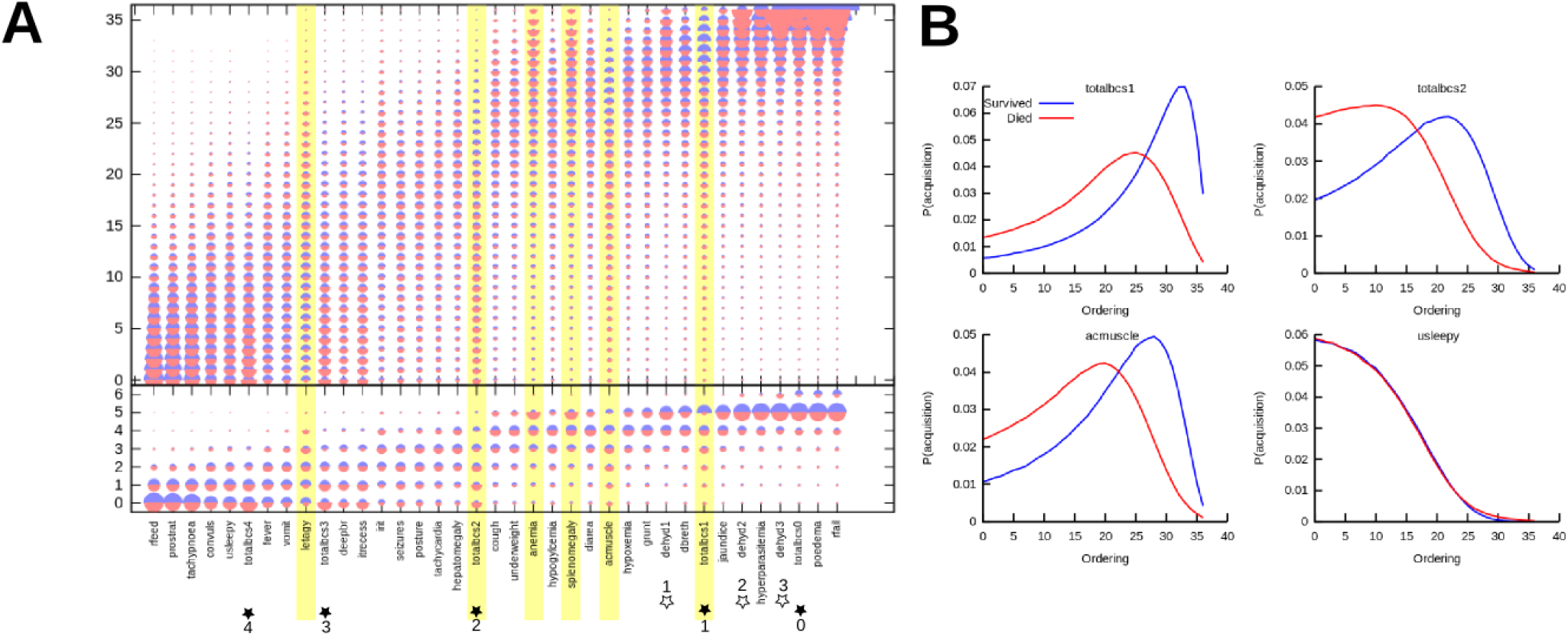
Inferring the pathways of malarial disease progression with HyperTraPS. (A) The HyperTraPS algorithm (see text) was used to infer the ordering with which malarial symptoms are likely acquired across patients. Horizontal axis records symptoms; vertical axis records ordering from low (early acquisition) to high (late acquisition). This ordering axis is grouped into 6 longer “ordering windows” in the lower subsection of the figure, to display broader trends in addition to specific features of the dynamics. The size of a semicircle denotes the posterior probability that a given symptom is acquired at a given ordering in progression of malaria. Red semicircles are posteriors from the dataset of patients who died; blue semicircles inferred from patients who lived. Highlighted symptoms display a greater KS distance between posteriors from survival and death pathways than between either posterior and the uninformative prior, forming potentially diagnostic features. (B) Posterior distributions on ordering for three features that differentiate between patients that eventually die and those that eventually survive, and for one that does not discriminate.

Intuitively, the inferred orderings for those features that are themselves progressive (decreasing *BCS scores* and increasing *dehydration indices*) match the expected disease course for an initially healthy patient whose condition worsens over time. Substantial separation of features is observed, with *refusal to feed* and *tachypnoea* among the earliest-onset features, and *renal failure* and *pulmonary oedema* among the latest. The posteriors for some features (*dehydration, BCS, refusal to feed, jaundice, hyperparasitaemia*) are more tightly constrained around given orderings than other features, suggesting varying flexibility amongst clinical features in the ordering in which they are manifested during disease progression.

Several clinical features notably discriminate the live and dead outcome groups. The dead cohort exhibits substantially earlier onset of *BCS scores*, consistent with the predictive power of these features identified in the above mutual information analysis. The dead cohort also shows later inferred ordering for *anaemia*, and earlier *accessory muscle use* and *deep breathing*. The posteriors for all these features were notably different between living and dead data classes, with greater Kolmogorov-Smirnov distances between posteriors from survival and death pathways than between either posterior and the uninformative prior (Fig 2C). The congruence between the features identified through this pathway analysis and those identified through the above mutual information approach suggest that this novel inference approach reveals robust aspects of disease progression, which we test further below.

### Validation of data-driven inference with independent expert survey

To validate our findings on disease progression, we surveyed 11 clinical practitioners in the field of malaria to build a consensus picture of clinical perceptions of malaria progression. We asked respondents to score each of the 25 features from our dataset as 1-2-3 (early; intermediate; late) according to their clinical perception of when a particular symptom was most likely to appear during disease progression. We then compared the mean response from the survey to the mean orderings inferred through HyperTraPS. We found a strong correlation between the clinician’s views and the orderings arising from computational inference from the full dataset (Fig 3, R = 0.518, p = 0.008). The views from the survey correlated better with inferred results from the subset of patients who lived (R = 0.521) than with those from patients who died (R = 0.399). The largest discrepancies between the inferred and survey orderings include fever, tachycardia, and coughing (which the survey ranks earlier than the inference) and prostration, convulsions, and coma (which the survey ranks later than the inference). The raw data for these features, in the form of frequency counts for each feature across patients, tend to support the inference picture: for example, prostration is seen to be an extremely common feature and coughing often occurs only in patients with several other symptoms (Fig 3).

**Figure 3.**
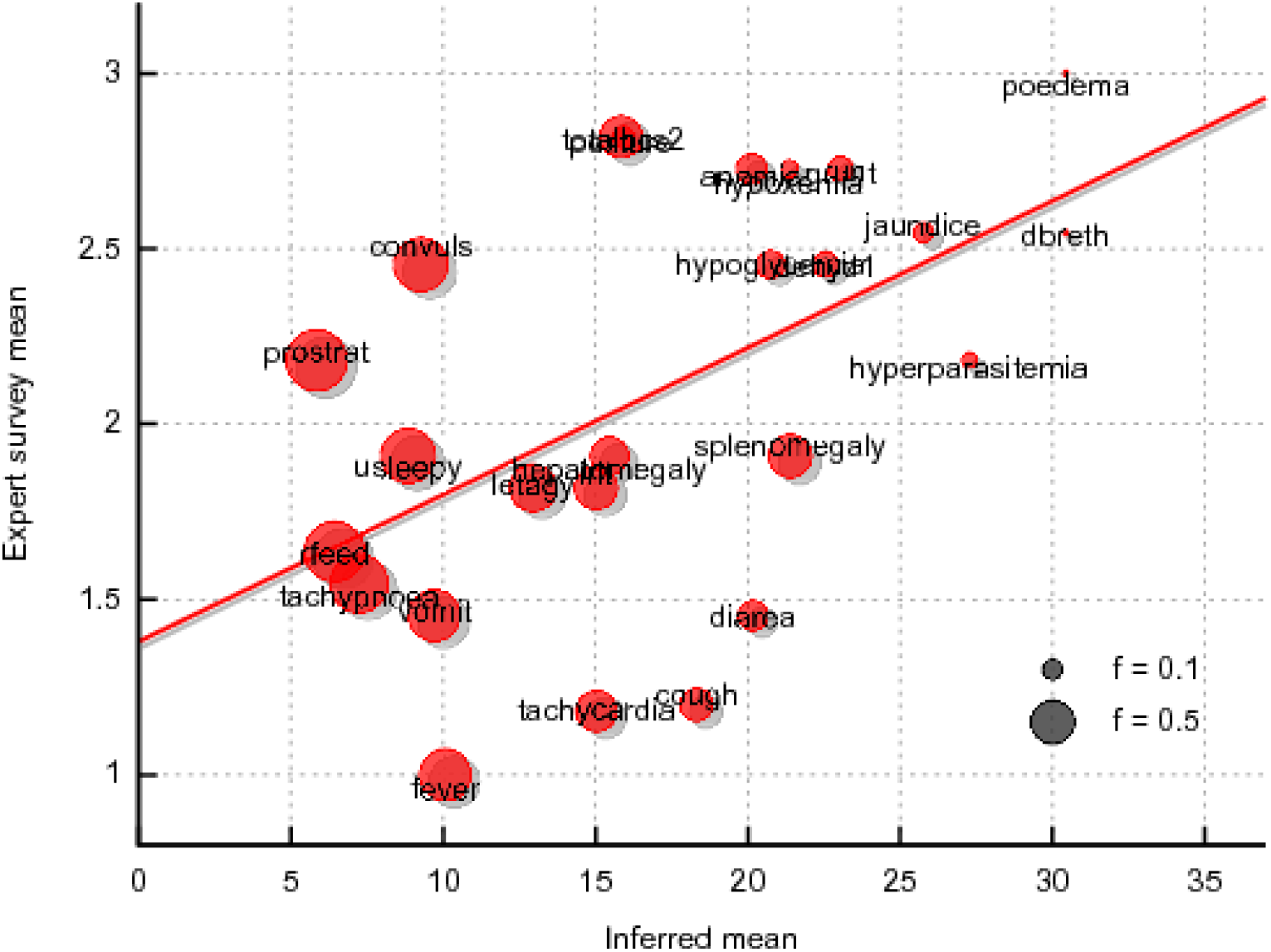
Comparison of machine learning results with expert survey. Horizontal axis gives the mean ordering of a symptom’s acquisition inferred by HyperTraPS; vertical axis gives the mean ordering of that symptom resulting from a survey of expert opinions (see text). The size of each circle represents the frequency *f* with which that feature is “present” when observed in the dataset: small circles are rarely observed, large circles commonly so.

### Using inferred pathway data to estimate hidden features

Our computed probabilistic description of the pathways underlying malaria progression allows us to predict unobserved features in particular patients. More precisely, by learning the structure of, and variability in, probabilistic pathways of disease progression, we can, in effect, build a set of probabilistic statements reporting the chance that an unobserved symptom is present or absent in a given patient, contingent on the presentation of other symptoms in light of the inferred ordering and relationship between symptoms. For example, the symptom `fever’ is inferred to occur consistently earlier than ‘hyperparasitaemia’; therefore, a patient whose *fever* status has not been observed, but who has been observed to present *hyperparasitaemia*, will be predicted to likely (but not certainly) have already acquired a fever. The pathways inferred by HyperTraPS generalise this simple two-feature picture to allow predictions based on the presence or absence of all features.

To demonstrate this predictive capability, we tested our algorithm on a randomly sampled subset of half of the patients from our dataset, learning the posterior distributions describing ordering of disease progression as in Fig. 4. We then artificially obscured 10% of the features in a random subset of 1,000 patients from the remaining (unseen) dataset and attempted to predict the values of these obscured features in different patients (Fig 4A). Of 3,318 features artificially obscured in the test dataset, we obtained predictions with 75% confidence for 1,330 features (40%). 83% of these predictions were correct, reflecting successes in predicting both presence and absence of obscured features. By comparison, a predictor only using the frequency counts of feature incidence in the dataset resulted in a 68% success rate using the same protocol.

**Figure 4.**
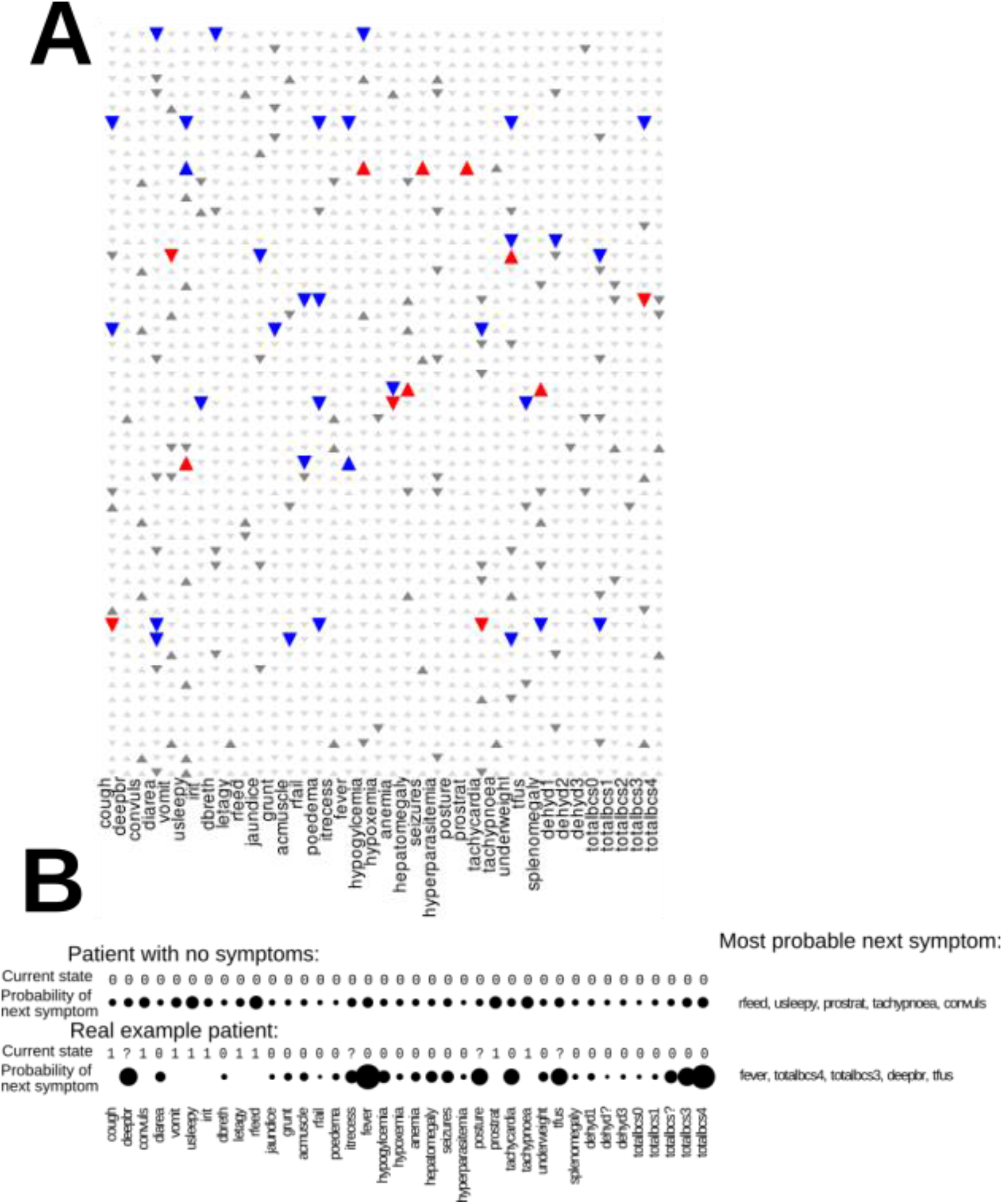
Prediction and validation of hidden patient symptoms using HyperTraPS. (A) Rows correspond to an illustrative subset of individual patients; columns give different observed symptoms. Upwards triangles denote feature presence, downwards triangles denote absence. A random subset of features were artificially hidden, and the prediction algorithm using HyperTraPS posteriors described in the text was then applied to predict the presence or absence of these features given the remaining features (small grey triangles). Blue triangles denote correct predictions; red denote incorrect predictions; large grey triangles give instance where no strong prediction was available. 83% of predictions (1104 of 1330) were accurate. (B) Illustration of prediction of likely next steps in disease progression for a given patient. Starting from any given patient state, HyperTraPS posteriors give the probability that any symptom is the next to be acquired by that patient. Circles represent the probability that each symptom will be acquired next, in two cases: a patient with no symptoms, illustrating the agreement with Fig 2, and a real patient taken from the dataset. In both cases the five most likely next symptoms are given.

### Using inferred pathway data to predict future progression

Another potentially more clinically valuable mode of prediction facilitated by our pathway analysis, is the likely next step in a disease progression pathway associated with an individual patient. Given an inferred set of likely progression pathways and a (possibly incomplete) observation of patient symptoms, we can integrate over all the possible states that could give rise to that observation and compute the probabilities with which each unacquired symptom will be the next step in a progression trajectory. Importantly, this approach may help target therapies to address the next most important stage in progression of the disease in individuals. To demonstrate this process, we used the learned pathways of disease progression to predict the likely next symptoms to become manifest as the disease progresses for a given sampled patient observation (Fig 4B). Due to the single-point nature of our dataset, we cannot use it to verify these predictions but we include this approach here as a demonstration for validation and a testable suite of hypotheses for future clinical studies.

### Using inferred pathway data to classify high-risk patients

HyperTraPS returns posterior distributions on the orderings of a progressive process, distinguishing events likely to occur earlier from those likely to occur later. The posteriors can be used to compute the probability that a patient with a given set of symptoms is on a high-risk disease progression pathway predicted to end in mortality, or on a lower-risk pathway predicted to end in survival.

To do so, we use Bayes’ theorem:

> P(survivor pathway | patient data) = P(patient data | survivor pathway) P(survivor pathway) / P(patient data)

to compute the likelihood associated with observing a given patient’s symptoms from the lower-risk (survivor) posterior parameterisations. There is an equivalent computation for the high-risk (dead) posterior (Fig 5A).

**Figure 5.**
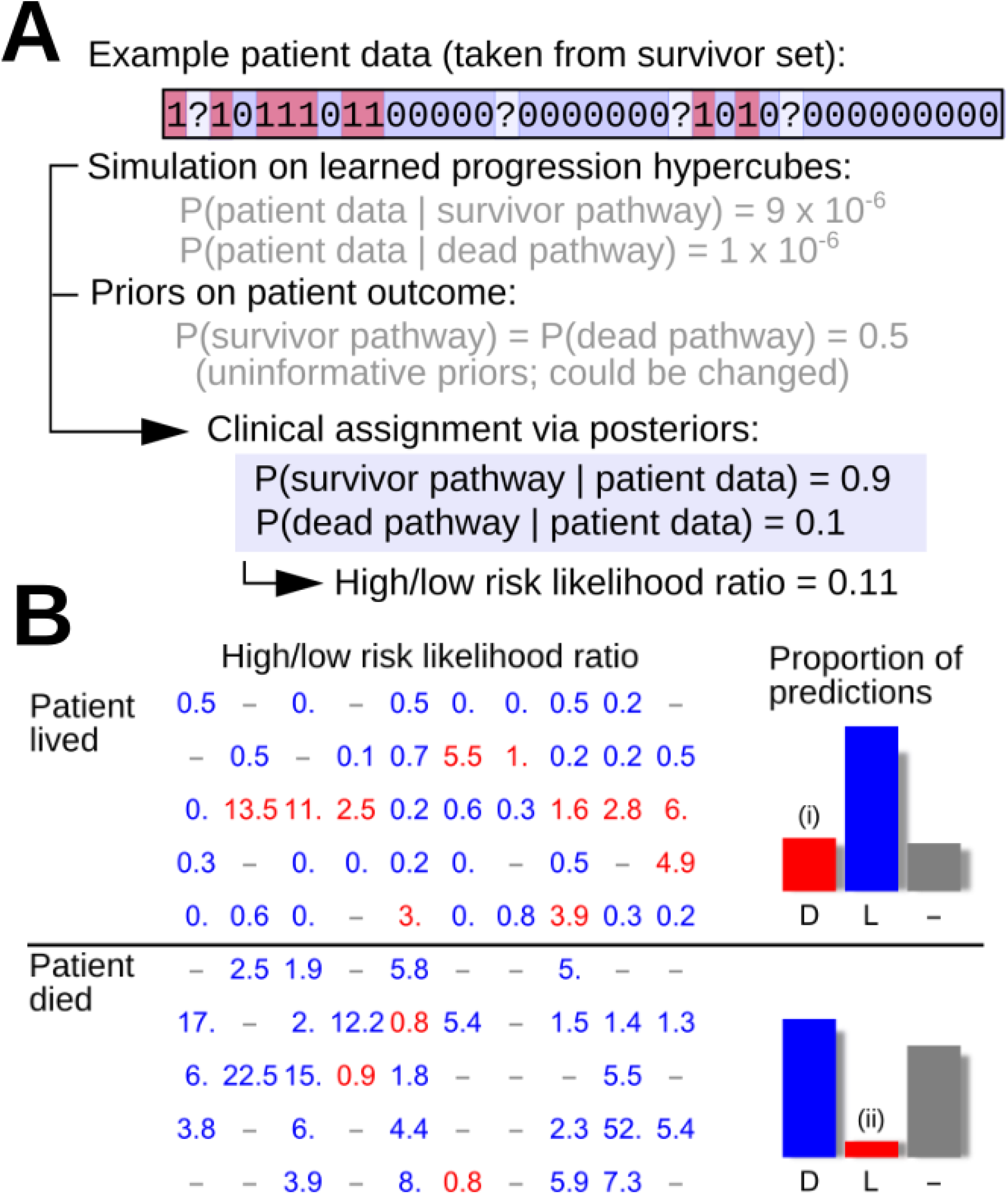
Bayesian classification of patient risk. (A) Applying Bayes’ theorem and simulation on the learned dynamics of surviving and dead patients to classify risk of new patients. (B) A test dataset of 50 patients that died and 50 patients that survived was analysed using posterior distributions for disease progression pathways derived from a separate training dataset. Figures give the likelihood ratio of a given patient being on a high-risk trajectory to that patient being on a low-risk trajectory, used to classify patients into high and low risk classes. Blue figures show where this classification aligns with the true patient outcome; red figures show where this classification does not align with patient outcome; dashes indicate cases where a classification was not available. Bars show the proportions of correct (blue) to incorrect (red) classifications. 81% of classifications (57 of 70) were successful; false positive identification rate of high-risk patients (i) is 20%, and false negative identification of high-risk patients (ii) is 6%.

We tested this approach by first assigning equal prior weight to both pathways (*P(survivor pathway) = P(dead pathway) = 0.5)* and then computing estimates of *P(patient data | survivor pathway)* and P(*patient data | dead pathway*) for 50 randomly chosen patients from each of the living and dead datasets (Fig 5B). If one predicted pathway was at least 1.5 times more likely than the other, we recorded this prediction as the more likely pathway for that patient. Of the 50 randomly chosen patients from the dead dataset, 23 had associated predictions: 22 were correctly predicted as high-risk and 1 was incorrectly predicted as low-risk. Of the 50 randomly chosen patients from the living dataset, 25 had associated predictions: 15 were correctly classified as low-risk and 10 were misclassified as high-risk. The risk of false negative high-risk classification from this approach is thus low, although a risk of false positive high-risk classification remains.

## Discussion

In this study, we have used mutual information and Bayesian inference of dynamic disease pathways with HyperTraPS to identify the most informative clinical features associated with death in Gambian children with severe malaria and the sequence of appearance of these features. This inference process reflects a novel way of analysing disease datasets, using cross-sectional data to learn the likely dynamics of disease progression. This is possible because the underlying model represents individual patient data as being sampled from that patient’s disease progression trajectory from healthy through the progressive acquisition of disease symptoms. By analysing many patient samples together and placing them all in the same probabilistic framework for disease progression, we can thus use a single-point observation to characterise the structure of, and variability in, progression pathways across a population. The value of this powerful approach is clear: we can simultaneously learn the dynamic pathways of disease progression, identify key predictors of clinical outcome, and use this unprecedented elucidation of disease dynamics to facilitate novel and clinically informative classification of the clinical risk associated with individual patients.

MI identified *Blantyre coma score* as the most informative clinical feature associated with death. A major prognostic feature of severe malaria is impaired consciousness, a feature which is adequately captured by the Blantyre Coma score (an ordinal score that ranges from 0 to 5, where a score of 5 indicates normal consciousness and a score of 0 indicates deep coma). We and others have previously reported the correlation of the coma score and the odds of death. A coma score of 2 or less defines cerebral malaria (CM) [1,5-7,12]. The absence of CM reduced the odds of death significantly. The next feature identified, respiratory distress (RD), increased the odds of death in patients with and without CM. In this model, both CM alone and CM in combination with RD accounted for the majority of cases with increased mortality. The administration of a blood transfusion was identified as the next informative prognostic feature in patients with CM and RD. Blood transfusion appeared to reduce mortality in those patients presenting with the three severe malaria syndromes (severe malarial anaemia [SMA], CM and RD). Interestingly, although the reduced odds of death were observed in patients with SMA, the majority of this group did not present with SMA. The potential benefit of transfusing patients with haemoglobin concentrations greater than 50 g/L is unclear and the WHO recommends blood transfusion for patients with haemoglobin concentrations up to 60 g/L only in presence of RD or impaired consciousness [13]. Our data suggest that patients with higher haemoglobin concentrations could benefit from blood transfusion and this seems particularly beneficial for patients presenting with CM and RD. Indeed, patients with CM and RD that received blood transfusions presented lower mortality than those not transfused, although their mean haemoglobin concentration (68.7 g/L) was above the recommended threshold for transfusion. Abnormal posturing is not frequently observed in non-comatose patients with SM. However, we identified abnormal posturing as an informative feature in patients with SMA and RD. A plausible explanation is that these patients presented with advanced RD and oxygen deprivation, exhibiting thus symptoms associated with hypoxic brain dysfunction.

We also observed that clinical features associated with a better clinical outcome (below average mortality), showed a good correlation with the WHO classification. In particular, we found that presence of an enlarged spleen (splenomegaly) was associated with a better outcome in patients with SMA. This is likely related to the spleen’s attempt to clear *P. falciparum* infected erythrocytes from blood, which probably reflects an adequate immune response.

In addition to identifying the most informative features that predict death, we used a Bayesian approach to infer the sequence of appearance of events that lead to death, providing the first posterior distributions on the progression dynamics of clinical symptoms in SM. This data-driven approach was validated by an independent survey of 11 experienced clinicians. Most clinicians who agreed to participate noted that they were uncertain about disease progression, as they commonly encountered only late-stage children in the clinical setting. There was general agreement among clinicians to score non-specific features such as fever, loss of appetite or prostration as early stage, whereas features of extreme severity, like deep coma, renal failure or jaundice were consistently placed as late features. Of note, some of the discrepancies observed between the clinical perception (survey) and the data-driven prediction can be reconciled. For example, the presence of fever or “reported fever” is scored as an early feature by clinicians since fever is the most likely guiding feature to suspect malaria. However, the data-driven algorithm may fail to capture this event since presence of fever is based on actual temperature recorded. Indeed, prior self-treatment with antipyretics (mostly paracetamol) can be as high as 50% [14]. On the other hand, neurological features such as coma and convulsions are commonly scored by clinicians as “late”, since these life-threatening features usually appear when the infection has not been identified and treated promptly [15]. Conversely, data-driven inference tends to score neurological features as “early” since the algorithm is blind to the time elapsed between the onset of symptoms and presentation to hospital. Also, this study was biased towards recruitment of more severe cases of *P. falciparum* malaria. An apparent limitation of the validation survey was that clinicians were asked to score features as early, intermediate or late. However, despite clinician uncertainty and the limited sample size of the survey, the level of agreement between expert clinical intuition and our prediction was remarkable.

Notably, we can use the differences in inferred dynamics of fatal and non-fatal SM cases to classify new patients according to their inferred risk. This novel approach captures not just a snapshot of individual risk factors but the full probabilistic information about the learned pathways of disease progression, allowing the histories of previous patients to inform the clinical analysis of new patients. This approach, validated with a test dataset, aligns with the goals of precision medicine and makes full use of available biomedical data. We indubitably anticipate its use in numerous other diseases and clinical contexts.

The relevance of the analysis we present in this research is twofold. Firstly, the prediction of clinical features associated with poor clinical outcome using mutual information validated prior findings and identified novel features. One of these features has potential translational impact and suggests the potential benefit of transfusing patients with higher haemoglobin concentrations beyond what is currently recommended by the WHO. These findings, however, must be validated in larger datasets and longitudinal studies. Secondly, the inferred sequence of events from a cross-sectional analysis is novel and important. For ethical reasons, clinical studies are characteristically not suited to study or describe the natural course of a disease, since treatment must be administered as soon as the diagnosis has been made and a treatment option is available. By inferring the sequence of events from cross-sectional data, this approach provides new insights into the natural history of disease *in the absence of longitudinal data*.

## Methods

### Study population

The study population consisted of 2,695 children aged 4 months to 15 years diagnosed with severe malaria according to the WHO definition. Children were admitted to the Royal Victoria Teaching Hospital (RVTH) from January 1997 to December 2009. The study was originally designed to study genetic variants associated with severe malaria [12]. The initial set of variables used for feature selection included those present in the case report form. The list of the variables included is described in supplementary Table 1. A detailed description of the study population and clinical features associated with death has been published elsewhere [13].

### Clinical definitions

Children aged 4 months to 15 years were eligible for enrolment if they had a blood smear positive for asexual *P. falciparum* parasites and met one or more WHO criteria for SM [14]): Coma (assessed by the Blantyre Coma Score [BCS][6]), severe anaemia (haemoglobin [Hb] < 50 g/L or packed cell volume [PCV]<15), respiratory distress or RD (costal indrawing, use of accessory muscles, nasal flaring, deep breathing), hypoglycaemia (<2.2 mM), decompensated shock (systolic blood pressure less than 70 mmHg), repeated convulsions (>3 during a 24 hour-period), acidosis (plasma bicarbonate <15 mmol/L) and hyperlactatemia (plasma lactate> 5 mmol/L). CM was defined as a BCS of 2 or less with any *P. falciparum* parasite density. SMA was defined as haemoglobin under 50 g/L. Hepatomegaly was defined as >2 cm of palpable liver below the right costal margin. Patients were enrolled in the study if informed consent was given by the parent or guardian. The study protocol was approved by the Joint Gambia Government / MRC Ethical Committee (protocol numbers 630 and 670).

### Laboratory measurements

Haemoglobin was measured with a haematology analyser (Coulter ® MD II, Coulter Corporation, USA), and parasite density was counted on Giemsa-stained thick and thin films.

### Data management and statistical analyses

The data were collected on standardized forms, double entered into a database and verified against the original.

### Mutual information analysis

#### Data curation

We removed 11 patients from the dataset whose clinical outcome (death) was not known and a further 21 patients for which more than half the features were unobserved or missing. The missing features of the remaining 2,883 patients were imputed using the k-nearest-neighbour algorithm. Firstly, we computed the Hamming distance between all pairs of patients, i.e. the distance between patients *i* and *j* is

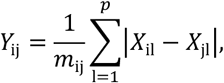

where *X_il_* denotes feature *l* of patient *i*, *p* is the number of features, and *m_ij_* is the number of features that are not missing for both patients. The clinical outcome was not included in the calculation of the distance matrix to avoid inducing artificial correlations between death and clinical features. Secondly, we set the missing features of patients equal to the median of the corresponding feature amongst the *k* nearest neighbours, i.e.

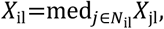

where *N_il_* is the set of *k* nearest neighbours of *i* for which feature *l* is not missing. We set *k = 13*, which is the square root of the number of complete cases [15]. Because *k* is odd, imputed values will also be binary.

#### Greedy, hierarchical partitioning of patients

We wanted to partition the patients into meaningful subsets to be able to predict clinical outcomes and identify different presentations of severe malaria. It was not feasible to consider a cross-tabulation of all available features because the number of cells would far exceed the number of patients. We thus considered a greedy algorithm that partitioned the patients into two subsets by the feature that was most predictive of death. The same step was recursively applied to the subsets until there were no more features that were predictive of death at a significance level *α = 0.05* after multiple hypothesis correction using the Holm-Bonferroni method [16].

We used mutual information (MI) between death and one of the features as a measure of predictive power because MI naturally quantifies the reduction in uncertainty about death achieved by observing said feature. MI has distinct advantages over the log odds ratio (LOR), which selects features with high sensitivity but low specificity: maximizing the LOR is likely to identify rare features that have a major impact on mortality, whereas maximizing MI will minimize uncertainty about death. We used the plugin estimator

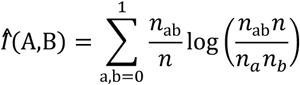

where *n* is the number of patients, *n_ab_* is the number of patients in the cell *ab* of the contingency table, and *n_a_*, *n_b_* are the number of patients in the a^th^ row and b^th^ column, respectively. This is a biased estimator [17]; correction strategies exist but we ignored the bias of the plugin estimator because we were more interested in which feature is most predictive than in the exact value of the MI.

We assessed statistical significance using a bootstrap algorithm. Firstly, we computed the unbiased estimates of the marginal distributions of the feature under consideration and mortality. Under the null hypothesis that the feature is not predictive of death, the joint distribution *q_ab_* is the product of the marginal distributions.

We drew 10^6^ independent samples of the same size as the dataset from the distribution under the null hypothesis and computed the MI for each synthetic dataset. The probability that the mutual information was larger than the observed value was equal to the proportion of simulated values that exceed the empirical one.

### HyperTraPS analyses

The HyperTraPS algorithm estimates the probability of observing a transition between two nodes *a* and *b* on a hypercubic transition graph, where edges are parameterised according to transition probabilities. The transitions we consider are between the state with no symptoms (assuming patients start healthy) and an observed set of symptoms in the dataset, corresponding to a particular time sample of a patient’s trajectory through the space of possible symptom patterns. We use an *LxL* matrix to encode the transition probabilities of the *L 2^L-1^* edges on the hypercube, assuming that each feature has a base rate of acquisition (*L* parameters) and may influence the rate of acquisition of all other features (*L(L-1)* parameters). We learned posterior distributions on these parameters by assigning them uninformative uniform priors and embedding HyperTraPS in an MCMC auxiliary pseudo-marginal (APM) algorithm [18]. Simulation of trajectories using parameterisations from these posteriors give then directly posterior distributions on orderings of feature acquisitions. To make predictions of unobserved features, we recorded points where these simulated trajectories matched the known features of a given sample and recorded the value(s) of the unobserved trait(s) at each of these points, building a tally of presence vs absence. To make predictions of future behaviour, we simply report the probabilities of subsequent steps from a given point on transition networks parameterised by these posteriors.

### Survey

The survey consisted of the question: “When do you expect to observe the following symptoms in the disease progression of malaria?” and the following features were listed: Prostration; Refusal to feed; Convulsions; Tachypnoea; Abnormal posturing (tonic seizures); Fever; Vomiting; Unusual sleepiness; Lethargy; Irritability; Coma (BCS <= 2); Tachycardia; Hepatomegaly (>2cm palpable liver); Coughing; Diarrhoea; Grunting; Anaemia (Haemoglobin < 50g/L); Splenomegaly (>2cm palpable spleen); Hypoglycaemia; Jaundice; Hyperparasitaemia (>250,000 / µL); Dehydration; Hypoxaemia (oxygen saturation <90%); Respiratory distress (intercostal recession, lower chest indrawing, use of accessory respiratory muscles, nasal flaring, deep breathing); Pulmonary oedema. Responses were returned between 16/01/2016 and 21/01/2016.

## Author Contributions

CCP and NJ had full access to all the data in the study and takes responsibility for the integrity of the data and the accuracy of the data analysis.

Study concept and design: TH, IJ, NJ, CCP

Acquisition of data: MJ, DK, CCP

Analysis and interpretation of data: TH, IJ, SG, OC, MB, NJ, CCP

Critical revision of the manuscript for important intellectual content: TH, IJ, MB, NJ, CCP Data analysis: TH, IJ, SG

Administrative, technical or material support: MJ, DK

Study supervision: IJ, NJ, CCP

## Conflict of Interest Disclosures

none declared

## Funding/Support

MalariaGEN’s primary funding is from the Wellcome Trust (grant number 077383/Z/05/Z) and from the Bill & Melinda Gates Foundation, through the Foundation for the National Institutes of Health (grant number 566) as part of the Grand Challenges in Global Health initiative. C.C-P is supported by the Medical Research Council (Clinician Scientist Fellowship: G0701885). IJ, SG, MB, NJ acknowledge grant EP/N014529/1

## Role of Sponsor

The WT and the MRC had no role in the design and conduct of the study.

## Acknowledgements

To the study participants and their parents/guardians. To the Royal Victoria Teaching Hospital (RVTH) nurses and fieldworkers: Yaya Dibba, Anthony Mendy and Abdoulie Camara; Senior laboratory technicians: Janet Riddle-Fullah andAbdou Bah; and Jalimory Njie (data entry clerk), Emmanuel Onykwelu (clinician), Augustine Ebonyi (clinician) and sister Haddy Njie. To MRC Laboratory technicians and assistants: Idrissa Sambou, Simon Correa, Madi Njie and Omar Janha. To Haddy Kanyi (data supervisor) and Mamkumba Sanneh (MRC Malaria Programme administrator).

## Supplementary Materials

**Table S1.** List of clinial features analysed.

| Feature | Coding | Definition | Missing |
| --- | --- | --- | --- |
| cough | binary | Coughing | 2.61% |
| hbtotal | numeric | Haemoglobin on admission derived from HB and PCV (g/dL) | 2.13% |
| totalbcs | ordinal | Total coma score on admission | 0.17% |
| deepbr | binary | Deep breathing | 64.77% |
| agemth | numeric | Age in months | 0.38% |
| convuls | binary | Reported convulsions prior to admission | 5.45% |
| coma | binary | Coma | 3.33% |
| comatime | numeric | Duration of coma in hours | 49.71% |
| diarea | binary | Diarrhea | 2.30% |
| vomit | binary | Vomiting | 2.06% |
| usleepy | binary | Unusually sleepy | 1.75% |
| irit | binary | Restless or irritable | 4.22% |
| dbreth | binary | Breathing difficulty | 2.81% |
| letagy | binary | Lethargy | 4.05% |
| rfeed | binary | Reduced feeding | 2.30% |
| pallor | 1 - conjunctiva,<br>2 - palms,<br>3 - tongue, 4 - all | Pallor (pale appearance) | 32.08% |
| jaundice | binary | Jaundice (yellowing of the skin) | 2.16% |
| grunt | binary | Grunting | 2.54% |
| acmuscle | binary | Use of accessory muscles of respiration | 2.26% |
| dehyd | 0 - not assessed, 1 -<br>nil, 2 - mild,<br>3 - moderate,<br>4 - severe | Dehydration | 21.17% |
| temp | numeric | Temperature (C) | 1.10% |
| hrate | numeric | Heart rate (bpm) | 0.31% |
| sao | numeric | Oxygen saturation | 66.07% |
| resprate | numeric | Respiratory rate (#/min) | 0.51% |
| spleen | numeric | Spleen size (cm) | 1.23% |
| liver | numeric | Liver size (cm) | 1.23% |
| rfail | binary | Renal (kidney) failure | 5.93% |
| poedema | binary | Pulmonary oedema | 5.93% |
| posture | 0 - not assessed,<br>1 - Yes, 2 - No | Abnormal posturing | 22.33% |
| prostrat | 0 - not assessed,<br>1 - Yes, 2 - No | Prostration (extreme weakness) | 23.26% |
| nofseizs | numeric | Number of seizures | 40.51% |
| waz | numeric | Weight z-score controlling for age | 29.43% |
| convadm | binary | Convulsions during admission | 65.42% |
| itrecess | binary | Intercostal recession | 70.46% |
| hypogl | binary | Hypoglycemia (glucose deficiency) | 29.95% |
| parmcl | numeric | Parasite count (#/μL) | 2.09% |
| death | binary | Death | 0.38% |
| cluster_idx | categorical | Cluster as inferred by the networks method | 0.00% |
| who_type | 1=CM, 2=CMRD,<br>3=RD, 4=<br>CMSMA,<br>5=CMRDSMA,<br>6=RDSMA,<br>7=SMA and<br>8=other | Malarial type (CM=cerebral malaria, RD=respiratory distress,<br>SMA=severe malarial anaemia) | 0.00% |

**Supplementary Figure 1.**
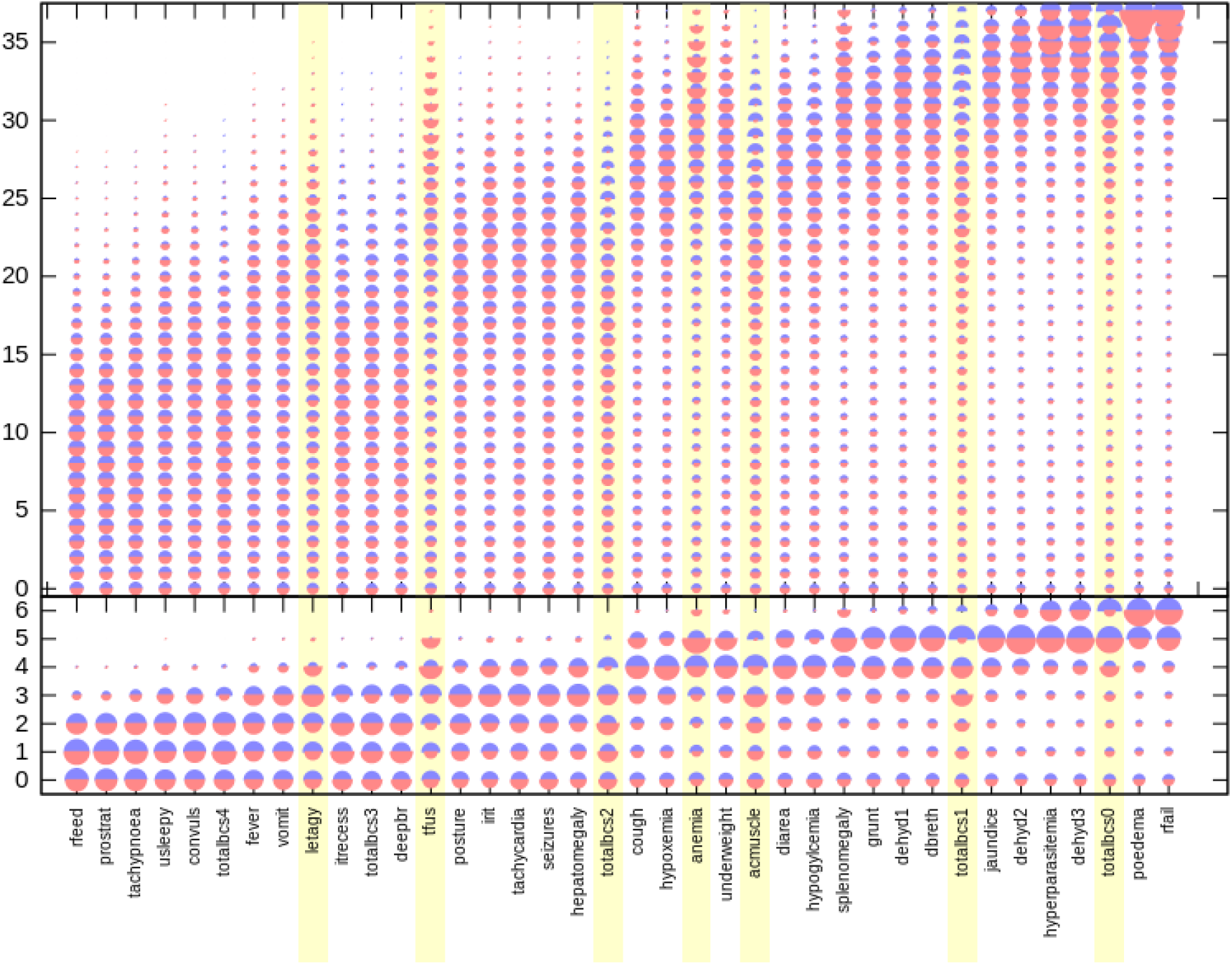
(related to Figure 2). Inferred disease progression pathways treating transfusion presence/absence as a disease feature. As in Figure 2, HyperTraPS was used to infer the ordering with which malarial symptoms are likely acquired across patients. Horizontal axis records symptoms; vertical axis records ordering from low (early acquisition) to high (late acquisition). The size of a semicircle denotes the posterior probability that a given symptom is acquired at a given ordering in progression of malaria. Red semicircles are posteriors from the dataset of patients who died; blue semicircles inferred from patients who lived. Highlighted symptoms display a greater KS distance between posteriors from survival and death pathways than between either posterior and the uninformative prior, forming potentially diagnostic features. When considered as a patient-intrinsic feature (though, in reality, an extrinsic intervention) transfusion is one of the strongest distinguishing features between the survivor and death pathway sets.

